# The distribution of mutational effects on fitness in *Caenorhabditis elegans* inferred from standing genetic variation

**DOI:** 10.1101/2020.10.26.355446

**Authors:** Kimberly J. Gilbert, Stefan Zdraljevic, Daniel E. Cook, Asher D. Cutter, Erik C. Andersen, Charles F. Baer

## Abstract

The distribution of fitness effects for new mutations is one of the most theoretically important but difficult to estimate properties in population genetics. A crucial challenge to inferring the distribution of fitness effects (DFE) from natural genetic variation is the sensitivity of the site frequency spectrum to factors like population size change, population substructure, and non-random mating. Although inference methods aim to control for population size changes, the influence of non-random mating remains incompletely understood, despite being a common feature of many species. We report the distribution of fitness effects estimated from 326 genomes of *Caenorhabditis elegans*, a nematode roundworm with a high rate of self-fertilization. We evaluate the robustness of DFE inferences using simulated data that mimics the genomic structure and reproductive life history of *C. elegans*. Our observations demonstrate how the combined influence of self-fertilization, genome structure, and natural selection can conspire to compromise estimates of the DFE from extant polymorphisms. These factors together tend to bias inferences towards weakly deleterious mutations, making it challenging to have full confidence in the inferred DFE of new mutations as deduced from standing genetic variation in species like *C. elegans*. Improved methods for inferring the distribution of fitness effects are needed to appropriately handle strong linked selection and selfing. These results highlight the importance of understanding the combined effects of processes that can bias our interpretations of evolution in natural populations.

## INTRODUCTION

Understanding the distribution of fitness effects (DFE) of new mutations is necessary to characterize the role of mutation in the evolutionary process and to determine the full impact that mutations have on the fitness of individuals and populations. The DFE influences the rate and trajectory of adaptive evolution (Orr 2000; Good et al., 2012), the maintenance of genetic variation (Charlesworth et al. 1995), the evolution of sex and recombination (Peck *et al.* 1997), the fate of small populations (Schultz and Lynch 1997), the molecular clock (Ohta 1992), the rate of decay of fitness due to Muller’s Ratchet (Loewe 2006), and the evolution of the mutation rate itself (Kondrashov 1995; Lynch 2008). Understanding the DFE is also necessary for accurate characterization of the genetic basis of complex traits, including human disease (Eyre-Walker 2010), and has been sought for decades (Boyko *et al.* 2008; Eyre-Walker and Keightley 2007; Kousathanas and Keightley 2013; Charlesworth 2015; Kim *et al.* 2017; Tataru *et al.* 2017). Any dynamic model of evolution must either define the DFE explicitly and assume its distribution or ignore the varying effects of mutations. Despite these fundamental roles, the DFE is a challenging property to estimate (Eyre-Walker and Keightley 2007; Charlesworth 2015).

The DFE defines the probability that a new mutation will alter organismal survival or reproduction by a given magnitude. Following common practice, we restrict our consideration of the DFE to new deleterious mutations that reduce fitness relative to the ancestral state. This DFE can be quantified in two basic ways: (1) from direct experimental measurement of fitness effects of new mutations or mutation panels (Thatcher et al. 1998; Sanjuán et al. 2004; DePristo et al. 2007) or (2) from indirect inference using the site frequency spectrum (SFS) of genetic variants in populations (Loewe *et al.* 2006; Keightley and Eyre-Walker 2007; Boyko *et al.* 2008). Using the SFS-based approach, population genomic data have been used to infer the DFE for many organisms (Boyko *et al.* 2008; Kousathanas and Keightley 2013; Charlesworth 2015; Kim *et al.* 2017; Tataru *et al.* 2017). With rare exceptions, these taxa are predominantly or obligately outcrossing. The SFS approach benefits from the large number of mutations, accessible with genome sequencing methods, that have experienced a fairly long evolutionary history in the natural environment. However, the utility of the SFS approach depends on the ability of analytical methods to use the observed standing genetic variation to correctly infer the effects of new mutations.

An accurate inference of the DFE allows one to understand the evolutionary trajectory that mutations will follow, because the selection coefficient conferred by any given mutation (*s*) combines with the effective size of the population in which it arose (*N*_e_) to determine the efficacy of selection (*N*_e_*s*) on that variant. The accuracy of the SFS inference method is challenged, however, by its sensitivity to non-equilibrium demography, population structure, and non-random mating (Eyre-Walker 2006). Demographic changes like population size expansions or contractions that mimic some effects of natural selection can lead to mischaracterization of the DFE (Eyre-Walker and Keightley 2007). Population size changes and cryptic population structure should largely be accounted for with existing methods that contrast two sets of loci: one set presumed to be selectively neutral (*e.g*., four-fold degenerate sites in coding sequences) and thus reflecting neutral demographic change, versus a second set presumed to experience the direct effects of selection (*e.g*., zero-fold degenerate sites in coding sequences) in addition to the neutral effects of demography (Keightley and Eyre-Walker 2007).

Non-random mating, of which self-fertilization is the most extreme form, presents a more difficult problem. Inbreeding reduces the effective recombination rate (Nordborg 1997), which exacerbates the effects of selection at linked loci (Cutter and Payseur 2013; Charlesworth and Wright 2001; Felsenstein 1974). With reduced recombination, individual variants may no longer have independent evolutionary trajectories, as the fitness effects of their nearby genomic neighbors can play a role in changing the allele frequency of a focal variant (Hill-Robertson interference; Hill and Robertson 1966). How Hill-Robertson interference affects inference of the DFE from the SFS cannot easily be predicted *a priori*. Furthermore, self-fertilization also exposes more variants as homozygous in the population, erasing possible effects of additivity in heterozygotes. In outcrossing taxa, selection against deleterious alleles typically occurs in heterozygotes, because recent deleterious mutations will be present as rare alleles that almost always occur in heterozygous state. Excess homozygosity caused by self-fertilization thus means that selection on homozygous genotypes will be a major driver of allele frequency change with potentially profound implications for inference of the DFE from variant frequencies. Few studies have inferred the DFE in non-obligately outcrossing organisms (Arunkumar *et al.* 2015; Huber *et al.* 2018), motivating deeper investigation into the impact of extreme selfing on DFE estimation.

Here, we report the DFE estimated from the SFS of a globally distributed collection of *Caenorhabditis elegans*, a nematode roundworm with a 99% to 99.9% rate of self-fertilization (Cutter *et al.* 2019). In this species, the rate of recombination varies along the holocentric chromosomes with low recombination in the central third of autosomes that also are gene-dense and enriched for essential genes (C. elegans Sequencing Consortium 1998; Rockman and Kruglyak 2009a; Cutter *et al.* 2009). These features stand in stark contrast to the genomes of many outcrossing taxa (*e.g*., *Drosophila* and mammals), in which regions of low recombination are depleted of genes, especially essential genes. In *C. elegans*, genome architecture combines with selfing to cause strong linked selection (Cutter and Payseur 2003; Crombie et al. 2019; Andersen *et al.* 2012a; Thomas *et al.* 2015). Selfing and linked selection in *C. elegans* contribute to a nearly 100-fold reduced polymorphism relative to the hyperdiversity of obligately outcrossing congeners (*e.g*., *C. remanei* and *C. brenneri*; (Cutter *et al.* 2013; Dey *et al.* 2013)). This influence also is observed within the genome: nucleotide diversity in low-recombination regions is five-fold lower than high-recombination regions (Rockman and Kruglyak 2009a; Andersen *et al.* 2012b; Thomas *et al.* 2015; Lee *et al.* 2020). We estimate the homozygous DFE for *C. elegans* by the maximum likelihood method implemented in the DFE-alpha software (Keightley and Eyre-Walker 2007; Eyre-Walker and Keightley 2009). Further, we evaluate the robustness of the inferred DFE from simulated data that matches the genome architecture and life history of *C. elegans* to understand how the joint effects of self-fertilization, genome structure, and natural selection influence estimates of the DFE.

## MATERIALS AND METHODS

### *C. elegans* genome-wide variant data

The *C. elegans* genome-wide variant data were acquired from the *C. elegans* Natural Diversity Resource 20200815 release (Cook *et al.* 2017). These data were generated by aligning Illumina short reads from each strain to the WS276 N2 reference genome (Lee *et al.* 2018) and then calling variants using the GATK4 (v4.1.4.0) *HaplotypeCaller* function and recalled using the *GenomicsDBImport* and *GenotypeGVCFs* functions (Poplin *et al.* 2018). Variants were annotated using SnpEff (v4.3.1) (Cingolani *et al.* 2012). To generate the site frequency spectra, we filtered the CeNDR VCF to contain the recently described 328 strain set (Lee *et al.* 2020), but two strains were removed. The strain ECA701 was removed because of high levels of residual heterozygosity, and the strain JU1580 was removed because it was found to be in the JU1793 isotype in the 20200815 CeNDR release. The final strain list contained 326 strains (Table S1).

The variants for all spectra were polarized using the XZ1516 strain as the ancestor, so this strain is not included in any spectra. For each SFS, we further pruned the VCF to contain only sites with no missing genotype data and with allele frequencies greater than 0%. We generated 18 SFS that encompass three different subsets of *C. elegans* strains, three different genomic regions, and two different site class comparisons. For the three different subsets of *C. elegans* strains, we used (1) the entire population sample (n = 325, 326 minus XZ1516), (2) the subset of “swept” strains (n = 273), and (3) the subset of “divergent” strains (n = 52). We classified a strain as “swept” if any of chromosomes I, IV, V, or X contained greater than or equal to 30% of the same haplotype (Andersen et al. 2012). Any strains not among the swept strains were classified as “divergent” (Table S1). For the three different genomic regions, we used (1) the whole genome, (2) high recombination chromosome arms, or (3) low recombination chromosome centers, as defined previously (Rockman and Kruglyak 2009). For the two different site class comparisons within coding sequences, we used (1) 4-fold vs 0-fold degenerate sites only or (2) 4-fold vs the combination of 0-fold degenerate sites and sites annotated as causing high or moderate deleterious effects as predicted by SnpEff (Cingolani et al. 2012). The R package SeqinR (Charif and Lobry 2007) was used to parse gene positions and a custom script was used to classify the degeneracy of each non-variant site as 0- or 4-fold degenerate. We counted the number of 0- and 4-fold degenerate sites that were invariant and included these sites as the 0% derived allele frequency class in each spectrum. We generated BED files (Quinlan and Hall 2010) of 0- and 4-fold degenerate sites using a custom AWK script that categorizes the alleles as ancestral or derived, into specific chromosomal regions, and by their predicted SnpEff effects. We then used vcfanno (Pedersen *et al.* 2016) to append these annotations to the original VCF file (Danecek *et al.* 2011). The annotated variant data was extracted from the VCF to a tab-separated file using BCFtools (Li 2011). This file was used to generate the 18 site-frequency spectra (File S1-S18).

### Simulation of site frequency spectra with SLiM

We used SLiM v2.1 (Haller and Messer 2019) to conduct forward-in-time simulations mimicking key features of *C. elegans* genome structure and life history. We simulated a single population with non-overlapping generations. Population size was constant in a given simulation with 99.9% of reproduction occurring via self-fertilization, as well as a comparison set of simulations with full outcrossing (*N* = 50,000 for outcrossing simulations and *N* = 500,000 in selfing populations; see Table S1). We generated a 24 Mb genome to represent the coding fraction of 100 Mb *C. elegans* genome, divided into six 4 Mb chromosomes comprising 1.44 Mb left and right arms and a 1.12 Mb center region (C. elegans Sequencing Consortium 1998). Recombination varied between the arms and centers of each chromosome: 2.35 × 10^−7^ crossovers per base pair per generation in arms, 4.96 × 10^−8^ crossovers per base pair per generation in centers (Rockman and Kruglyak 2009b). We simulated mutations to arise at a uniform rate across the genome (3.3 × 10^−9^ mutations per base pair per generation, or approximately 0.08 mutations per simulated genome per generation), with 75% of mutations subject to selection and the remaining 25% neutral to match the incidence of replacement- and synonymous sites in coding sequences (Saxena *et al.* 2019). Of the 75% of mutations with fitness effects, simulation sets created either all selected mutations as deleterious or as 99.9% deleterious plus 0.1% beneficial. Beneficial mutations had a gamma-distributed DFE with mean selection coefficient *s* = 0.01, shape parameter *β* = 0.3, and additive effects (*h* = 0.5). The deleterious mutational effects followed a mixture of two gamma distributions to best match the hypothesis for biologically realistic DFEs. Most of this mixture distribution (95%) was defined by a gamma distribution made up of many nearly neutral deleterious mutations with mean *s* = −0.001, shape *β* = 0.3, and dominance, *h* = 0.3. The remaining 5% of the distribution of deleterious fitness effects came from a gamma distribution comprising mutations with strong and more recessive effects to imitate the existence of recessive lethals with mean *s* = −0.01, *β* = 0.3, and *h* = 0.2. We also conducted a second set of simulations with a more extreme DFE for deleterious mutations: a single gamma distribution with *s* = −0.161, *β* = 2.13, and *h* = 0.3 to serve as an example from a more severely deleterious distribution of mutations, similar to that inferred from the empirical *C. elegans* dataset.

### DFE inference with DFE-alpha

We used DFE-alpha (Keightley and Eyre-Walker 2007) to infer the DFE by maximum likelihood from both the empirical and simulated datasets. We then compared the DFE derived from the empirical and simulated datasets to test for deviations between the inferred DFEs that might result from inaccurate or oversimplified genomic architectures and evolutionary parameters.

We then conducted a second set of simulations that used DFE parameters inferred from the empirical data as input for the simulations, re-estimated the simulated DFE, and tested whether we could accurately recover this underlying DFE. We also compared the DFE inferred for polymorphisms linked to different chromosome regions (arms versus centers). We used DFE-alpha with the two-epoch model and the folded site frequency spectrum, as recommended by the DFE-alpha documentation and also as is commonly used across empirical applications of the method. We also averaged the site frequency spectrum across three sampling points in the simulations, at generations 4*N,* 4.5*N*, and 5*N* prior to applying the inference approach as performed previously (Messer and Petrov 2013).

### Data availability

The authors state that all data necessary for confirming the conclusions presented in the article are represented fully within the article. All data and methods required to confirm the conclusions of this work are within the article, figures, and supplemental materials. SFS data from *C. elegans* strains is additionally archived on GitHub at: https://github.com/Thatguy027/SFS_Invariant_Sites/tree/master/2020_SFS_Analysis/Paper_Files/manuscript_spectra

## RESULTS

We first used DFE-alpha to estimate the distribution of deleterious fitness effects from the genomes of 325 strains of *C. elegans*. Across all subsets of the data, the inferred distributions show the highest densities for strongly deleterious mutations, with mutations of weakest effect being second most prevalent. For analyses using only the 4- and 0-fold degenerate sites designated as neutral or selected, respectively, 62.5% of mutations fall into the most severely deleterious class and 17.3% into the weakest, nearly neutral deleterious class (whole genome; Figure 1A). Chromosome arms and centers have qualitatively similar profiles, but with arms exhibiting a somewhat greater density of highly deleterious mutations (Figure 1). We also inferred the DFE using a more sophisticated categorization of selected sites beyond simply 0-fold degenerate positions of coding sequences. In this approach, we predicted deleterious functional effects of variants (SnpEff, see Methods), including stop-gained, splice-site, and non-synonymous variants. The overall profile for the DFE in these cases exhibited decreased weight in the strongest deleterious class (49.7% of mutations for the whole genome) at the expense of more sites of weakly deleterious effects being inferred (23.2% for the whole genome; Figure 1B). Again, chromosome arms showed a greater relative weight in the most deleterious class compared to chromosome centers.

**Figure 1.**
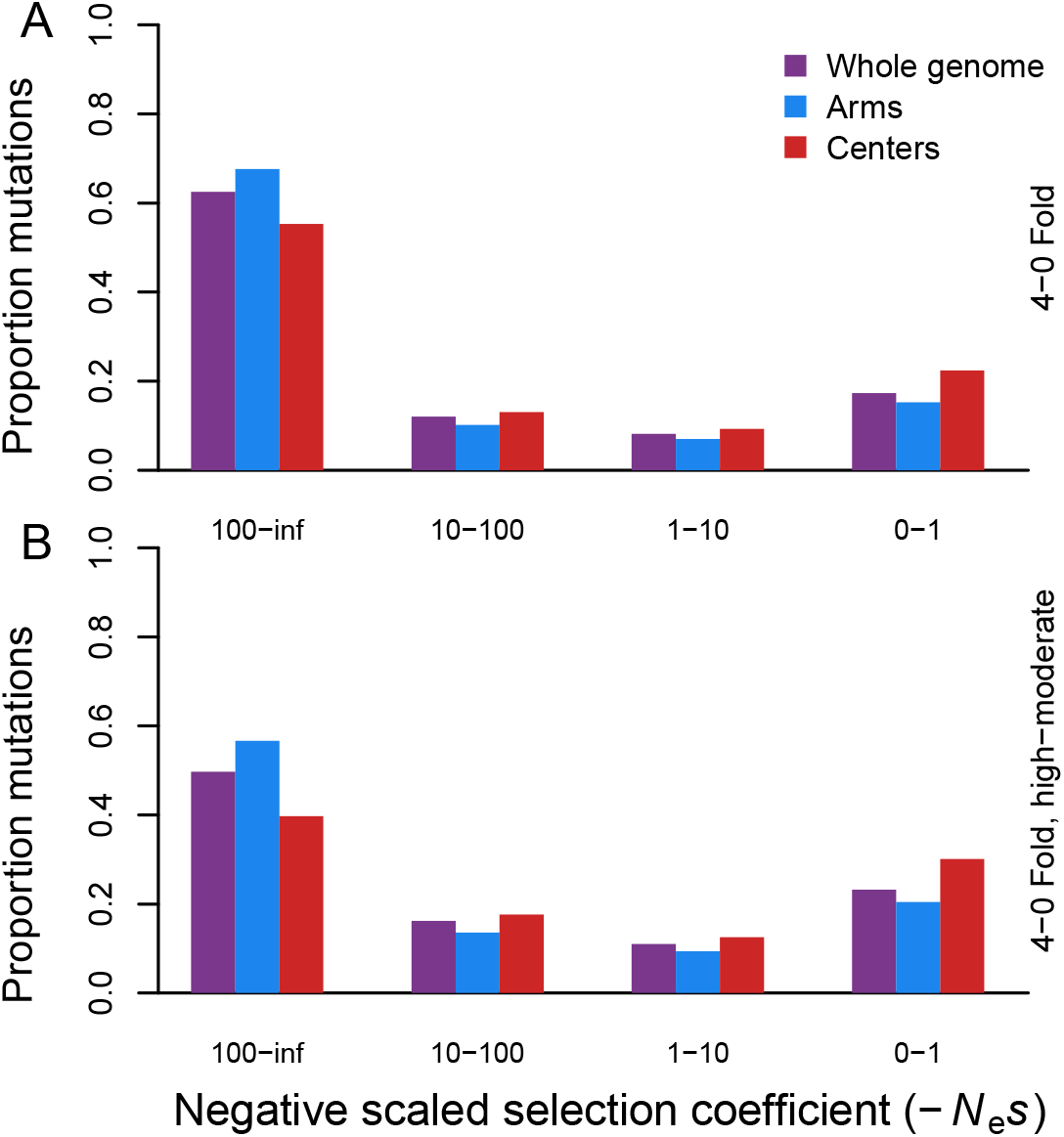
DFE inference for *C. elegans* based on coding sequence polymorphism from genomes of 325 strains. Site frequency spectra of polymorphisms derive from the whole genome (purple) or separately for coding sequences in chromosome arms (blue) and centers (red), defined by recombination rate boundaries (Rockman and Kruglyak 2009a). The neutral and selected classes for site frequency spectra provided to DFE-alpha correspond to 4-fold degenerate sites and 0-fold degenerate sites, respectively (A). An alternative selected site class also included the sites characterized as having variants exerting high or moderate fitness effects by SnpEff (B). All estimated parameters of the inferred DFE are listed in Table 1.

To assess the reliability of these DFE inferences from standing variation under extreme selfing, we conducted a series of forward-time simulations that mimic *C. elegans* genome architecture and reproduction. Inference of the DFE from simulated data sets, where the true input DFE is known, resulted in different inferred estimates than expected from the input mixture gamma distribution (Figure 2A; Supplementary Table 2). This inferred distribution comprised almost entirely nearly neutral mutations (−*N*_e_*s* = 0-1), regardless of linkage to chromosome arms or centers (Figure 2A). These results also differed starkly from what we observed for the empirical *C. elegans* results. Our second set of simulations used a more strongly deleterious input DFE, similar to, but more extreme than, the DFE inferred from the *C. elegans* data. In our simulations with high selfing, DFE-alpha was better able to estimate the input DFE for this mutational spectrum that was weighted toward strongly deleterious mutations (Figure 2B). Curiously, however, it showed a U-shaped distribution with excess density in the nearly neutral class relative to the input (Figure 2B), reminiscent of the pattern observed in the empirical analysis of *C. elegans* genomes (Figure 1).

**Table 1.** Inferred parameters of the DFE for the empirical *C. elegans* dataset using DFE-alpha. The mean selection coefficient (Es) is not scaled by effective population size (*i.e.*, not *N*_e_*s*). Mean (absolute) selection coefficients greater than 1 reflect the long tail of a leptokurtic gamma distribution. See Keightley and Eyre-Walker 2007 for explanation and further interpretation.

| Dataset | Genome portion | SNP set | Population size after first epoch, in 2-epoch mode ($N_2$, relative to $N_1 = 100$) | Duration of epoch after first population size change ($t_2$) | Weighted recent effective population size ($N_w$, Keightley & Eyre-Walker 2009) | Mean selection coefficient ($E_s$) | Shape parameter ($b$) |
| --- | --- | --- | --- | --- | --- | --- | --- |
| Empirical full strain set | Whole | 4-0-fold | 1000 | 299.23 | 225.06 | -40.42 | 0.17 |
|  | Arms |  | 1000 | 245.70 | 204.04 | -124.88 | 0.16 |
|  | Centers |  | 1000 | 298.62 | 224.83 | -22.36 | 0.15 |
|  | Whole | 4-0-fold, high-moderate | 1000 | 299.23 | 225.06 | -6.80 | 0.17 |
|  | Arms |  | 1000 | 245.70 | 204.04 | -20.83 | 0.16 |
|  | Centers |  | 1000 | 298.62 | 224.83 | -2.95 | 0.15 |
| Empirical swept strain set | Whole | 4-0-fold | 1000 | 679.00 | 359.09 | -6.63 | 0.20 |
|  | Arms |  | 1000 | 628.18 | 342.59 | -197.65 | 0.14 |
|  | Centers |  | 1000 | 501.51 | 299.61 | -0.72 | 0.26 |
|  | Whole | 4-0-fold, high-moderate | 1000 | 679.00 | 359.09 | -1.45 | 0.20 |
|  | Arms |  | 1000 | 628.18 | 342.59 | -29.71 | 0.13 |
|  | Centers |  | 1000 | 501.51 | 299.61 | -0.18 | 0.28 |
| Empirical diverged strain set | Whole | 4-0-fold | 307 | 140.07 | 142.22 | -546.09 | 0.13 |
|  | Arms |  | 372 | 143.18 | 147.62 | -321.05 | 0.15 |
|  | Centers |  | 1000 | 1814.82 | 636.79 | -16.75 | 0.14 |
|  | Whole | 4-0-fold, high-moderate | 307 | 140.07 | 142.22 | -71.53 | 0.13 |
|  | Arms |  | 372 | 143.18 | 147.62 | -50.33 | 0.15 |
|  | Centers |  | 1000 | 1814.82 | 636.79 | -2.37 | 0.13 |

**Figure 2.**
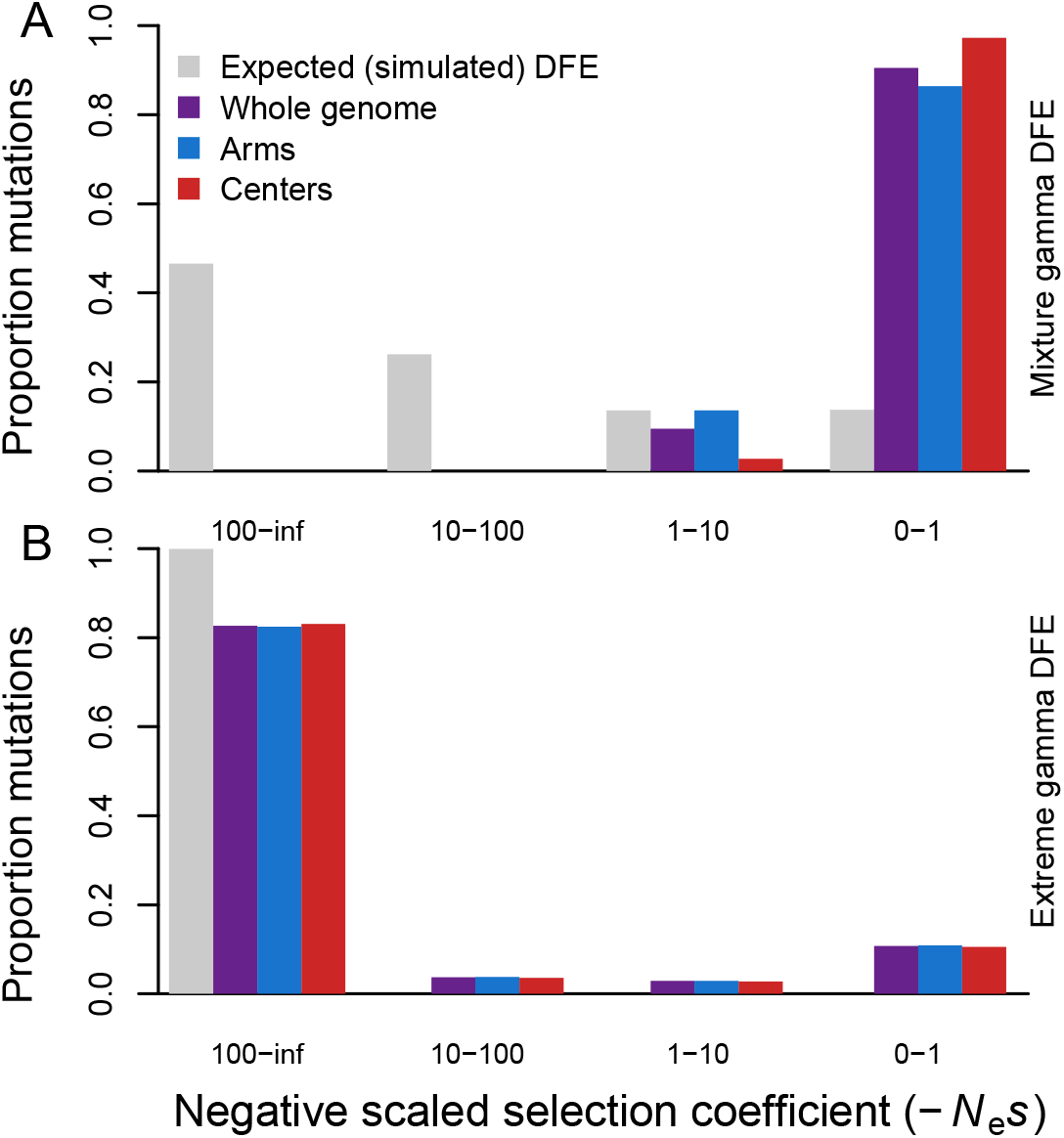
Distributions of deleterious fitness effects for simulations mimicking *C. elegans* genome structure and selfing reproductive mode. (A) The inferred DFEs for deleterious mutations (colored bars) from simulations with an input DFE (gray bars) of a mixture gamma distribution (β = 0.3 for 95% mutations with mean *s* = −0.001 and *h* = 0.3 plus 5% with mean *s* = −0.01 and *h* = 0.2). (B) Inferred DFEs for a more extreme, deleterious gamma distribution of mutational input (β = 2.13, mean *s* = −0.161, *h* = 0.3), similar to that of the inferred DFE from the empirical data. Larger values of −*N*_e_*s* are more deleterious; simulation census size *N* = 500,000; selfing rate 99.9%; SFS are averaged over generations 4*N,* 4.5*N*, and 5*N*. All estimated parameters of the inferred DFE are listed in Table S2.

To further elucidate the impact and potential biases introduced by selfing on our DFE inferences, we contrasted simulations that differed only in reproductive mode: 99.9% selfing (Figure 2) versus 100% outcrossing (Figure 3). We found that the inferred DFE much more accurately matches the known input DFE for the simulations under a regime of full outcrossing (Figure 3) as opposed to high selfing. Both the mixture gamma DFE and the extreme DFE that we simulated matched well to the input mutational spectra under a regime of outcrossing, compared to the extremely poor match to the mixture DFE for highly selfing populations (Figure 2 *cf*. Figure 3). Nevertheless, the gene dense and low recombining chromosome centers exhibited an inferred DFE shifted toward weaker fitness effects even in these purely outcrossing simulations.

**Fig 3.**
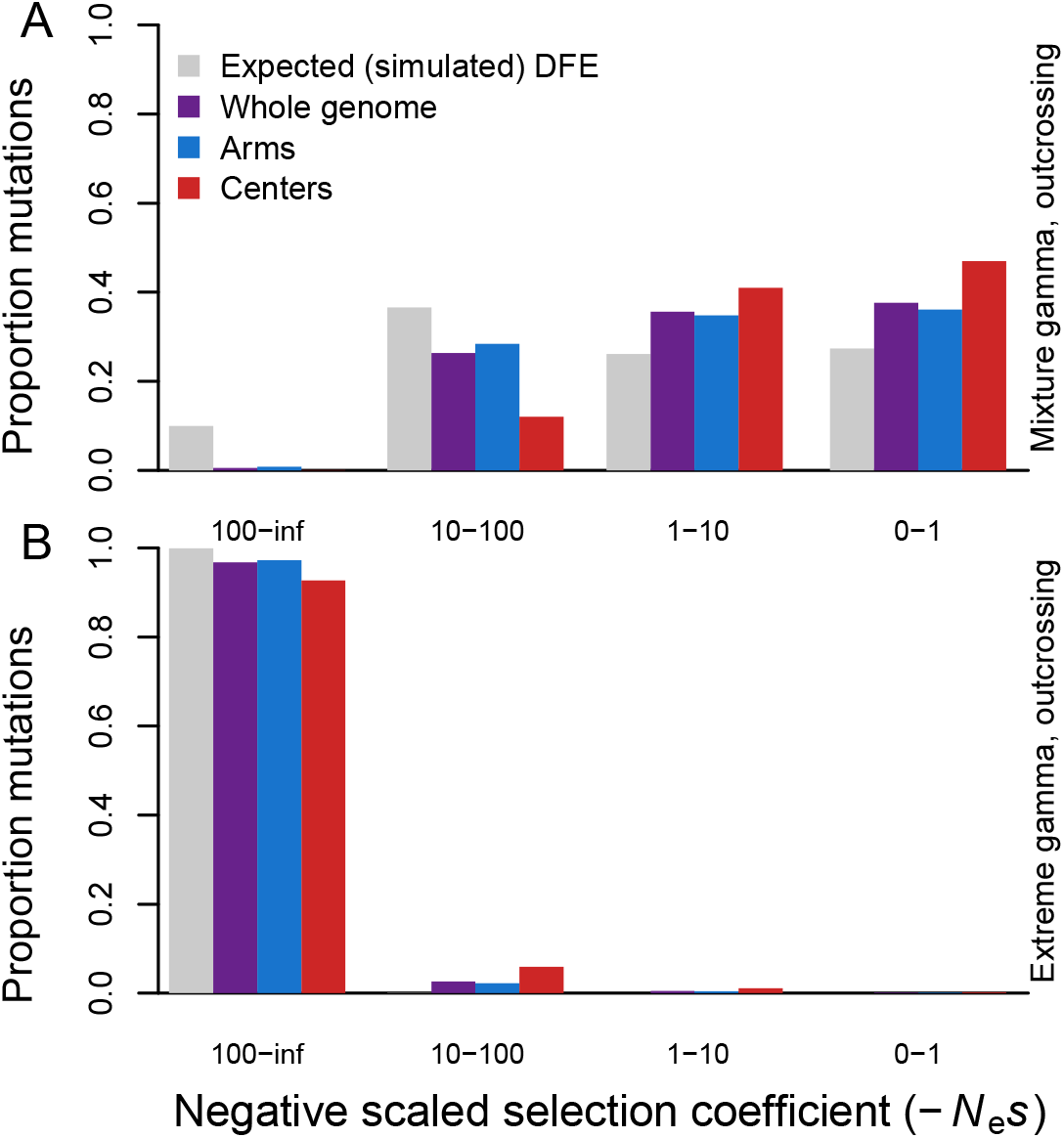
DFE inferences from simulations of fully outcrossing individuals. Genetic properties are otherwise equivalent to the selfing simulations shown in Figure 2, but with census size *N* = 50,000 instead of 500,000 (hence the difference in expectation of the gray bars from Figure 2). Panel (A) shows results of simulations using the mixture gamma distribution while panel (B) corresponds to simulations of an input DFE that includes greater incidence of highly deleterious mutations. SFS are averaged over generations 4*N,* 4.5*N*, and 5*N*. All estimated parameters of the inferred DFE are listed in Table S2.

We hypothesized that a subset of *C. elegans* strains might contribute to a perturbed DFE because they are hypothesized to have experienced selective sweeps that impacted large portions of the genome (Andersen et al. 2012). When we compared the DFE inferred using data from the 273 “swept” strain subset to the DFE inferred using data from the 52 non-swept “divergent” strain subset, we did not observe a drastic difference in DFE shapes for the whole genome analyses (Figure 4). Qualitatively, both subsets of the data showed the largest proportion of deleterious mutations in the strongest selection class and the second most abundant mutations to be in the weakest selection class (Figure 4). For chromosome centers, however, the swept strain subset showed a less strongly “U-shaped” distribution, instead having the second most weight for intermediate effect deleterious mutations (−*N*_e_*s* = 10-100; Figure 4B). Using the SFSs that included sites specified by SNPeff further exacerbated the DFE shift toward weaker fitness effects in chromosome centers for swept strains (Figure 4C,D). This pattern of weaker mutational effects inferred for the strains and genomic regions most impacted by selective sweeps suggests that linked selection influences the DFE inference.

**Figure 4.**
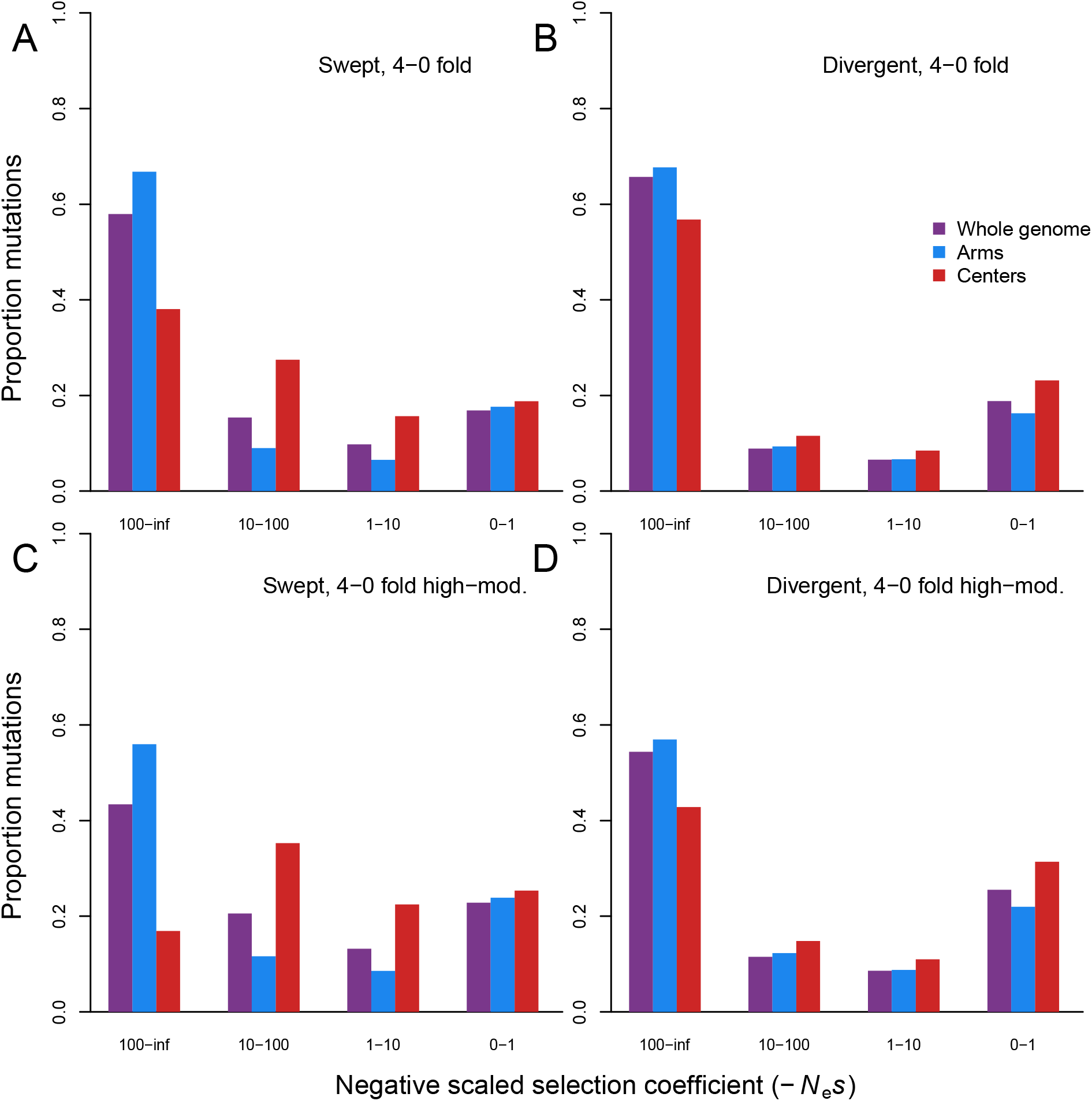
DFE inferences from subsets of *C. elegans* strains. (A, C) DFEs based on 273 “swept” strains with >30% of the genome showing strong influence of linked selection (Andersen et al. 2012). (B, D) DFEs based on 52 “divergent” strains with no evidence of selective sweeps across their genomes (see Methods). A and B show inferences that used only 0-versus 4-fold sites whereas C and D include sites designated as exerting high or moderate effects by SnpEff in the selected class. All estimated parameters of the inferred DFE are listed in Table 1.

We also hypothesized that the presence or absence of beneficial mutational input might complicate inference of the DFE in the context of selfing-induced homozygosity and linked selection because DFE-alpha only infers the DFE for deleterious mutations. Therefore, we tested whether the inclusion or exclusion of beneficial mutations in the simulations impacted the inferred DFE under high selfing. We observed that the presence of beneficial mutations most strongly influenced the fraction of mutations in the weak and intermediate fitness effect classes (−*N*_e_*s* = 0-1 and 1-10), shifting the inferred DFE toward a greater density of weakly deleterious effects relative to simulations that lacked any beneficial mutations (Figure 5). The pattern for chromosome centers contrasted starkly with chromosome arms and the whole genome when only deleterious mutations were present, exhibiting a more even distribution across all greater deleterious effect classes but still retaining a large proportion of mutations in the nearly neutral class (Figure 5B). Finally, we tested whether DFE inferences using the unfolded SFS might better match the expected DFE, but found no clear improvement relative to the performance of DFE-alpha using the folded SFS (Figure S1).

**Figure 5.**
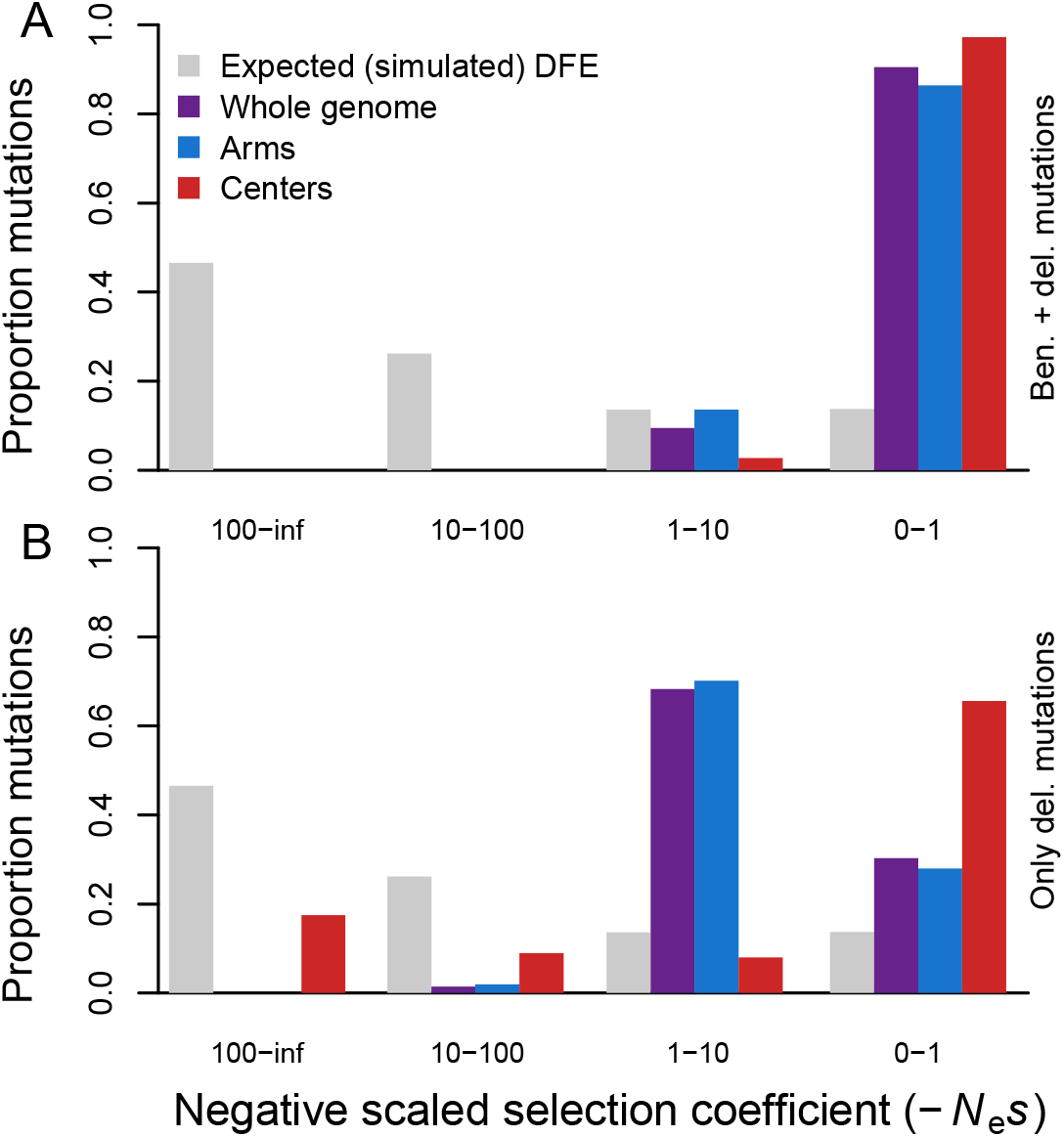
The DFE inferred from simulations with both deleterious and beneficial mutations (A, as in Figure 2A), or with deleterious mutations only (B). Simulation parameters as in Figure 2. SFS are averaged over generations 4*N,* 4.5*N*, and 5*N*. All estimated parameters of the inferred DFE are listed in Table S2.

## DISCUSSION

Population genomic data, in principle, provide a rich collection of allelic variation from which to infer the distribution of fitness effects (DFE) for mutational input into populations. Accurate estimation of the DFE from such data can be complicated by population demography that differs from equilibrium, due to growth or decline in population size, although methods implemented in inference algorithms attempt to minimize such demographic biases (Keightley and Eyre-Walker 2007; Boyko et al. 2008; Tataru et al. 2017). The influence of non-random mating on DFE inference, however, remains incompletely understood. Our exploration of the DFE for deleterious mutations in *C. elegans*, using genome sequences for 326 wild isolates combined with biologically motivated simulations, demonstrates that extreme self-fertilization can compromise accurate inference of the DFE.

Based on naive application of DFE-alpha to infer *C. elegans*’ DFE, one would conclude that mutational effects exert predominantly strong deleterious consequences on fitness. However, we showed that, when simulating conditions that mimic *C. elegans*’ genome architecture and its extreme 99.9% selfing mode of reproduction, the same inference algorithm does not always recover the input DFE. This discrepancy is not a general problem with the method, as simulations under random mating or less extreme selfing do successfully recover the input DFE. This contrast highlights the skew that a highly selfing mating regime can cause for inferred DFEs. Recent work in self-compatible *Eichhornia* found that the DFE could be recovered adequately with 98% selfing (Arunkumar *et al.* 2015), implying that the breakdown of the DFE inference arises toward the extreme of selfing seen in *C. elegans*. Using an alternative inference method, previous results indicated that a scaled additive model could be applied for the case of 97% selfing in *Arabidopsis* (Huber et al. 2018), though our results again suggest that such an assumption may break down with more extreme selfing. Both of these studies in plants benefited from calibrating their analysis of selfing populations with a close outgroup that is fully outcrossing. Although *C. elegans* lacks a known outgroup that would be appropriate to use in this way, future analysis of the DFE for the selfing *C. briggsae* in combination with its close outcrossing relative *C. nigoni* is promising for this comparative approach.

Our simulations show that the extreme selfing of *C. elegans* interacts with the shape of the DFE such that some distributions show a greater disparity between the DFE input by mutation and the output DFE inferred from polymorphism. This discrepancy depends on at least two factors: the nature of the underlying DFE (weighted to a more negative extreme versus weighted to a more nearly neutral distribution), and linkage between beneficial and deleterious variants. Dominance plays little role in the evolutionary fate of mutations in highly selfing populations, because new mutations become homozygous after only a few generations. As a consequence, selfing is thought to more effectively purge large-effect deleterious recessive mutations as compared to nearly additive weak-effect mutations (Charlesworth and Charlesworth 1998). This differential purging might induce the DFE inferred from polymorphism data to show an abundance of variants with nearly neutral deleterious effects, as we have seen for our simulations of high selfing leading to poor recovery of the underlying DFE when that distribution contained mutations over a wide range of effects (Figure 2A). Only under a simulated DFE where the vast majority of mutations entering the population are highly deleterious does the DFE inference appropriately recover the underlying distribution (Figure 2B), likely because few or no weakly deleterious mutations are segregating in the population at all. Indeed, our selfing simulations and *C. elegans* analysis both show high densities of mutations in either the highest or the lowest deleterious fitness class of mutations and few mutations of intermediate effect (Figure 1, Figure 2).

The effect of linkage between selected variants on the DFE inference is detectable from both our empirical and simulation datasets. Comparison of our “swept” versus “divergent” empirical data subsets reveals a pattern of weaker mutational effects inferred for the strains and genomic regions most impacted by selective sweeps (*i.e*., chromosome centers in swept strains). Considering chromosome arm and center regions separately also reflects the stark difference in recombination rate and gene density found between these regions in the *C. elegans* genome (C. elegans Sequencing Consortium 1998; Cutter et al. 2009; Rockman and Kruglyak 2009). This chromosomal heterogeneity leads to profound effects of linked selection in chromosome centers (Cutter and Payseur 2003; Andersen et al. 2012; Crombie et al. 2019), which we hypothesized might influence DFE inference. Indeed, we found that the inferred DFE tended to shift towards weaker-effect mutations in chromosome centers. To some extent, this effect was apparent even in simulated genomes with full outcrossing. However, we observed the most substantial differences between arm and center regions in our analysis of the subset of *C. elegans* data from strains that show selective sweeps that span nearly two-thirds of the genome. DFE-alpha can therefore account for modest perturbations due to linked selection within its demographic correction by approximating this influence as a reduction in effective population size (Keightley and Eyre-Walker 2007). The magnitude of effects from linked selection in *C. elegans* (Andersen et al. 2012; Thomas et al. 2015; Cutter and Payseur 2003), however, appear too extreme to be adequately accounted for in this way.

Furthermore, inclusion or exclusion of beneficial mutations in our simulations also greatly impacted the inferred DFE. The distribution inferred in all of our scenarios by DFE-alpha is that of deleterious mutations only, yet for the same input deleterious DFE, we can observe different inferences based on the presence or absence of beneficial mutations entering the population. In the presence of linkage between deleterious and beneficial variants, the inferred DFE for deleterious mutational effects is shifted to a more weakly deleterious distribution overall (Figure 5A cf. 5B). The difference in inference between simulated arms and centers of the genome also clearly shows this pattern of a greater weight for weakly deleterious mutations in the lower-recombining regions of the chromosome centers.

We conclude that the disparities between mutational input and the DFE inferred from site frequency spectra are caused by linked selection that is exacerbated by selfing. The consequences of linked selection are even further exacerbated by *C. elegans’* genome structure with gene-dense and low recombination chromosome centers. The impact of linked selection on the DFE inference was not due to underlying mutational differences along chromosomes, as centers and arms did not differ in selective effects of mutations in our simulations. The correction for demographic change implemented in DFE-alpha does not seem able to accommodate the degree of linked selection resulting from the extreme selfing experienced by *C. elegans*, since this correction only modulates the effective population size in an attempt to accommodate the presence of non-standard population conditions. The high values of N2 (the effective population size during the second epoch estimated by DFE-alpha and used for scaling the selection coefficient; see Table 1) capped at 1000 estimated during the DFE inference may reflect this poor fit to the demographic model due to selfing and linked recombination and thus the poor inferences of the DFE. Such an inability to fully account for the effects of strong linked selection conforms with the fact that the influence of selection on a genomic region is imperfectly approximated by a scaled effective population size (Kaiser and Charlesworth 2009; Neher 2013). Similar to Messer and Petrov (2013), we find that this bias causes genomic regions most impacted by linked selection to show an inferred DFE of weaker fitness effects. This problem is likely to be especially acute for populations that experience high rates of selfing.

As we strive to understand more about the role of deleterious mutations in evolution and the prevalence and distribution of their fitness effects, inferring the DFE in long-standing model systems is an essential first step towards a comprehensive understanding of the mutational process and the impacts of selection, demography, and genomic architecture on the fate of new mutations. We emphasize the difficulties that can be encountered when applying existing methods for inferring the DFE to non-standard population conditions, in particular for the extreme of non-random mating reflected by the high selfing of *C. elegans*. This challenge highlights the need for integration of empirical and theoretical approaches, and new methods, to account for perturbing effects of linked selection and non-random mating to generate a fuller understanding of the mutational processes that underlie the evolution of populations in the wild.

## ACKNOWLEDGEMENTS

KJG was funded by EMBO long-term fellowship ALTF2-2016 and Swiss National Science Foundation Ambizione grant #PZ00P3_185952. ADC is supported by a Discovery Grant from the Natural Sciences and Engineering Research Council (NSERC) of Canada. CFB and ECA were supported by NIH award R01GM107227.

## SUPPLEMENTARY FIGURES

**Figure S1.**
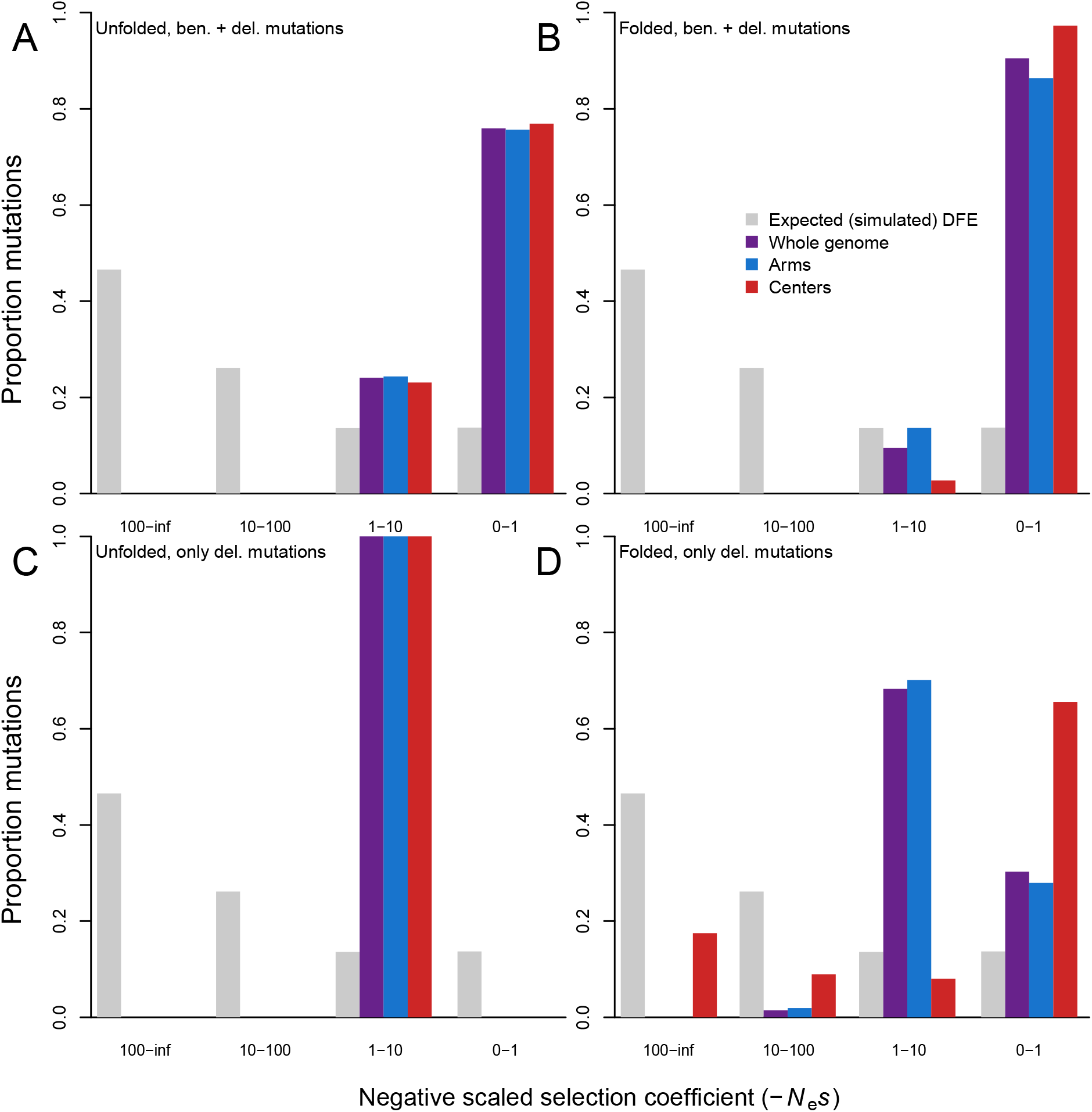
DFE-alpha analysis of simulated datasets using unfolded SFS as input (A, C) versus the folded SFS as input (B,D, as in Figure 5). These analyses performed more poorly than folded analyses at matching the input DFE (gray bars), particularly when simulations only included deleterious mutations (C,D), so therefore were not used in further analyses. All estimated parameters of the inferred DFE are listed in Table S2.

**Table S1.** The set of 326 *C. elegans* strains used in this study, and their classification as “swept” or “divergent” (see main text Methods for details).

| strain | strain_set | strain | strain_set | strain | strain_set | strain | strain_set |
| --- | --- | --- | --- | --- | --- | --- | --- |
| AB1 | swept | EG4946 | swept | JU3134 | swept | NIC268 | divergent |
| BRC20067 | swept | GXW1 | swept | JU3135 | swept | NIC269 | swept |
| BRC20263 | swept | JT11398 | swept | JU3137 | swept | NIC271 | swept |
| CB4851 | swept | JU1088 | swept | JU3140 | swept | NIC272 | swept |
| CB4852 | swept | JU1172 | swept | JU3141 | swept | NIC274 | swept |
| CB4853 | swept | JU1200 | swept | JU3144 | swept | NIC275 | swept |
| CB4854 | swept | JU1212 | swept | JU3166 | swept | NIC276 | swept |
| CB4855 | swept | JU1213 | swept | JU3167 | divergent | NIC277 | swept |
| CB4856 | divergent | JU1242 | swept | JU3169 | swept | NIC3 | swept |
| CB4857 | swept | JU1246 | swept | JU3224 | swept | NIC501 | swept |
| CB4858 | swept | JU1249 | swept | JU3225 | swept | NIC511 | swept |
| CB4932 | swept | JU1395 | swept | JU3226 | divergent | NIC513 | swept |
| CX11254 | swept | JU1400 | swept | JU3228 | swept | NIC514 | swept |
| CX11262 | swept | JU1409 | swept | JU323 | swept | NIC515 | swept |
| CX11264 | swept | JU1440 | swept | JU3280 | swept | NIC522 | swept |
| CX11271 | swept | JU1491 | swept | JU3282 | swept | NIC523 | swept |
| CX11276 | swept | JU1530 | swept | JU3291 | swept | NIC526 | swept |
| CX11285 | swept | JU1543 | swept | JU346 | swept | NIC527 | swept |
| CX11292 | swept | JU1568 | swept | JU360 | swept | NIC528 | swept |
| CX11307 | swept | JU1581 | swept | JU367 | swept | NIC529 | swept |
| CX11314 | swept | JU1586 | swept | JU393 | swept | PB303 | swept |
| CX11315 | swept | JU1652 | swept | JU394 | swept | PB306 | swept |
| DL200 | swept | JU1666 | swept | JU397 | swept | PS2025 | swept |
| DL226 | swept | JU1792 | swept | JU406 | swept | PX179 | swept |
| DL238 | divergent | JU1793 | swept | JU440 | swept | QG2075 | swept |
| ECA189 | divergent | JU1808 | swept | JU561 | swept | QG2811 | swept |
| ECA191 | divergent | JU1896 | swept | JU642 | swept | QG2813 | swept |
| ECA347 | divergent | JU1934 | swept | JU751 | swept | QG2818 | swept |
| ECA348 | swept | JU2001 | swept | JU774 | swept | QG2823 | swept |
| ECA349 | swept | JU2007 | swept | JU775 | swept | QG2824 | swept |
| ECA36 | divergent | JU2016 | swept | JU778 | swept | QG2825 | swept |
| ECA363 | divergent | JU2017 | swept | JU782 | swept | QG2827 | swept |
| ECA369 | divergent | JU2106 | swept | JU792 | swept | QG2828 | swept |
| ECA372 | divergent | JU2131 | swept | JU830 | swept | QG2832 | swept |
| ECA396 | divergent | JU2141 | swept | JU847 | swept | QG2835 | swept |
| ECA592 | swept | JU2234 | swept | KR314 | swept | QG2836 | swept |
| ECA593 | divergent | JU2250 | swept | LKC34 | swept | QG2837 | swept |
| ECA594 | swept | JU2257 | swept | MY1 | swept | QG2838 | swept |
| ECA640 | swept | JU2316 | divergent | MY10 | swept | QG2841 | swept |
| ECA703 | divergent | JU2464 | swept | MY16 | divergent | QG2843 | swept |
| ECA705 | divergent | JU2466 | swept | MY18 | swept | QG2846 | swept |
| ECA706 | divergent | JU2478 | swept | MY2147 | swept | QG2850 | swept |
| ECA710 | divergent | JU2513 | swept | MY2212 | swept | QG2854 | swept |
| ECA712 | divergent | JU2519 | swept | MY23 | divergent | QG2855 | swept |
| ECA722 | divergent | JU2522 | swept | MY2453 | swept | QG2857 | swept |
| ECA723 | divergent | JU2526 | divergent | MY2530 | swept | QG2873 | swept |
| ECA724 | divergent | JU2534 | swept | MY2535 | swept | QG2874 | swept |
| ECA730 | divergent | JU2565 | swept | MY2573 | swept | QG2875 | swept |
| ECA732 | divergent | JU2566 | swept | MY2585 | swept | QG2877 | swept |
| ECA733 | divergent | JU2570 | swept | MY2693 | swept | QG2932 | swept |
| ECA738 | divergent | JU2572 | swept | MY2713 | swept | QG536 | swept |
| ECA740 | divergent | JU2575 | swept | MY2741 | swept | QG556 | swept |
| ECA741 | divergent | JU2576 | swept | MY518 | swept | QG557 | swept |
| ECA742 | divergent | JU2578 | swept | MY679 | swept | QW947 | swept |
| ECA743 | divergent | JU258 | swept | MY772 | swept | QX1211 | divergent |
| ECA744 | divergent | JU2581 | swept | MY795 | swept | QX1212 | swept |
| ECA745 | divergent | JU2586 | swept | MY920 | swept | QX1233 | swept |
| ECA746 | divergent | JU2587 | swept | N2 | swept | QX1791 | divergent |
| ECA760 | divergent | JU2592 | swept | NIC1 | swept | QX1792 | swept |
| ECA768 | divergent | JU2593 | swept | NIC1049 | swept | QX1793 | divergent |
| ECA777 | divergent | JU2600 | swept | NIC1107 | swept | QX1794 | divergent |
| ECA778 | divergent | JU2610 | swept | NIC1119 | swept | RC301 | swept |
| ECA807 | divergent | JU2619 | swept | NIC166 | swept | WN2001 | swept |
| ECA812 | divergent | JU2800 | swept | NIC195 | swept | WN2002 | swept |
| ECA923 | swept | JU2811 | swept | NIC199 | swept | WN2033 | swept |
| ECA928 | swept | JU2825 | swept | NIC2 | swept | WN2050 | swept |
| ECA930 | swept | JU2829 | swept | NIC207 | swept | WN2063 | swept |
| ED3005 | swept | JU2838 | swept | NIC231 | swept | WN2064 | swept |
| ED3011 | swept | JU2841 | swept | NIC236 | swept | WN2066 | swept |
| ED3012 | swept | JU2853 | swept | NIC242 | swept | XZ1513 | divergent |
| ED3017 | swept | JU2862 | swept | NIC251 | divergent | XZ1514 | divergent |
| ED3040 | swept | JU2866 | swept | NIC252 | swept | XZ1515 | swept |
| ED3046 | swept | JU2878 | swept | NIC255 | swept | XZ1516 | ancestor |
| ED3048 | swept | JU2879 | swept | NIC256 | swept | XZ1672 | swept |
| ED3049 | swept | JU2906 | swept | NIC258 | swept | XZ1734 | swept |
| ED3052 | swept | JU2907 | swept | NIC259 | swept | XZ1735 | swept |
| ED3073 | swept | JU310 | swept | NIC260 | swept | XZ1756 | swept |
| ED3077 | swept | JU311 | swept | NIC261 | swept | XZ2018 | swept |
| EG4347 | swept | JU3125 | swept | NIC262 | swept | XZ2019 | divergent |
| EG4349 | swept | JU3127 | swept | NIC265 | divergent | XZ2020 | swept |
| EG4724 | swept | JU3128 | swept | NIC266 | swept |  |  |
| EG4725 | swept | JU3132 | swept | NIC267 | swept |  |  |

**Table S2.**
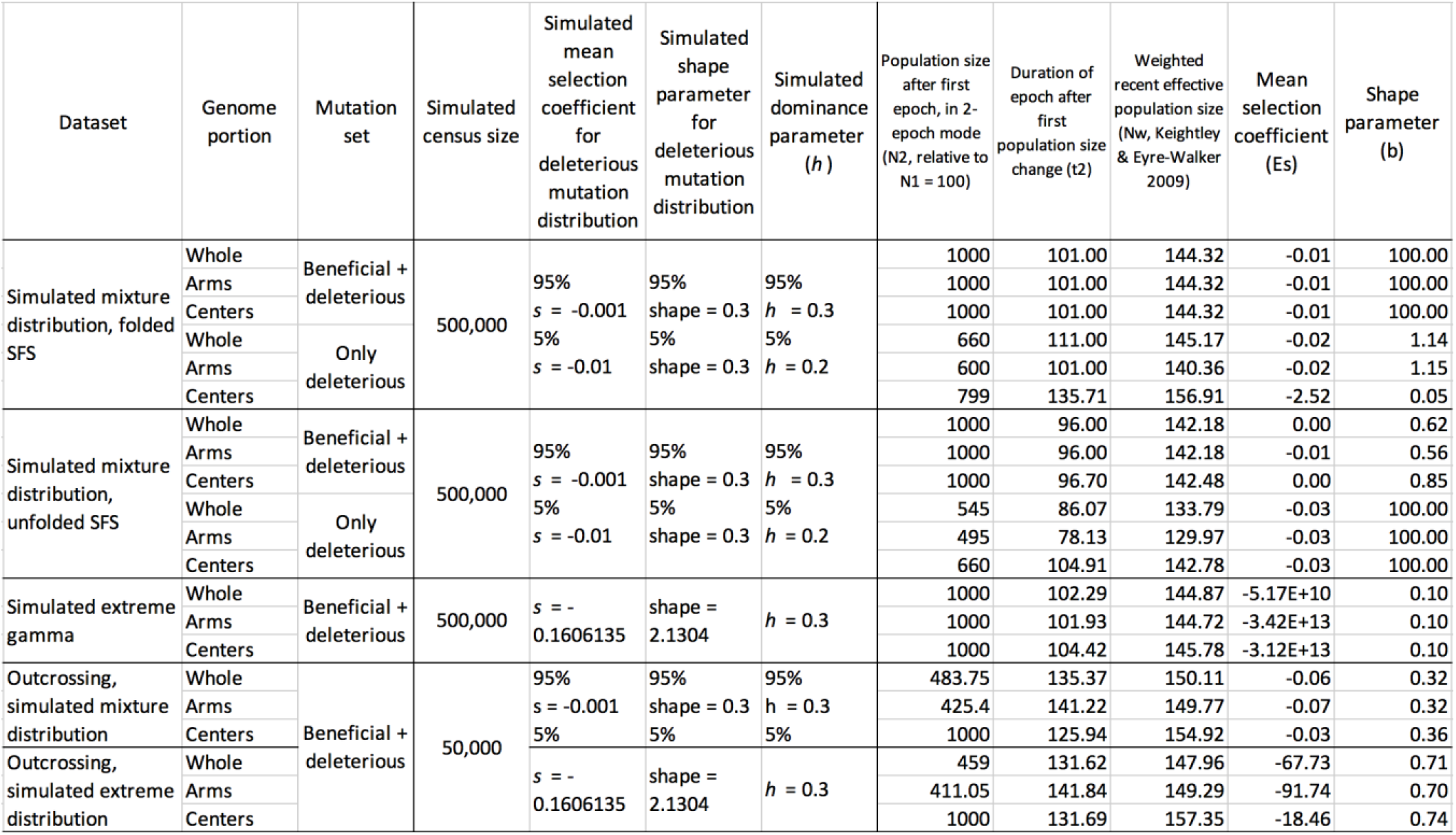
Inferred parameters of the DFE for the simulated datasets using DFE-alpha, averaged over 20 replicate simulations in each row.

## LITERATURE CITED

Andersen, E. C., J. P. Gerke, J. A. Shapiro, J. R. Crissman, R. Ghosh, et al., 2012a Chromosome-scale selective sweeps shape Caenorhabditis elegans genomic diversity. Nat. Genet. 44: 285–290. https://doi.org/10.1038/ng.1050

Andersen, E. C., J. S. Bloom, L. Kruglyak, M.-A. Félix, R. Ghosh, et al., 2012b Chromosome-scale selective sweeps shape *Caenorhabditis elegans* genomic diversity. Nat. Genet. 44: 285–290. https://doi.org/10.1038/ng.1050

Arunkumar, R., R. W. Ness, S. I. Wright, and S. C. H. Barrett, 2015 The evolution of selfing is accompanied by reduced efficacy of selection and purging of deleterious mutations. Genetics 199: 817–829. https://doi.org/10.1534/genetics.114.172809

Boyko, A. R., S. H. Williamson, A. R. Indap, J. D. Degenhardt, R. D. Hernandez, et al., 2008 Assessing the evolutionary impact of amino acid mutations in the human genome. PLoS Genet. 4: e1000083. https://doi.org/10.1371/journal.pgen.1000083

C. elegans Sequencing Consortium, 1998 Genome sequence of the nematode C. elegans: a platform for investigating biology. Science 282: 2012–2018. https://doi.org/10.1126/science.282.5396.2012

Charif, D., and J. R. Lobry, 2007 SeqinR 1.0-2: A Contributed Package to the R Project for Statistical Computing Devoted to Biological Sequences Retrieval and Analysis, pp. 207–232 in Structural Approaches to Sequence Evolution: Molecules, Networks, Populations, edited by Bastolla U., Porto M., Roman H. E., Vendruscolo M. Springer Berlin Heidelberg, Berlin, Heidelberg.

Charlesworth, D., and S. I. Wright, 2001 Breeding systems and genome evolution. Current Opinion in Genetics & Development 11: 685–690.

Charlesworth, B., 2015 Causes of natural variation in fitness: evidence from studies of Drosophila populations. Proc. Natl. Acad. Sci. U. S. A. 112: 1662–1669. https://doi.org/10.1073/pnas.1423275112

Cingolani, P., A. Platts, L. L. Wang, M. Coon, T. Nguyen, et al., 2012 A program for annotating and predicting the effects of single nucleotide polymorphisms, SnpEff: SNPs in the genome of Drosophila melanogaster strain w1118; iso-2; iso-3. Fly 6: 80–92. https://doi.org/10.4161/fly.19695

Cook, D. E., S. Zdraljevic, J. P. Roberts, and E. C. Andersen, 2017 CeNDR, the Caenorhabditis elegans natural diversity resource. Nucleic Acids Res. 45: D650–D657. https://doi.org/10.1093/nar/gkw893

Cutter, A. D., A. Dey, and R. L. Murray, 2009 Evolution of the Caenorhabditis elegans genome. Mol. Biol. Evol. 26: 1199–1234. https://doi.org/10.1093/molbev/msp048

Cutter, A. D., R. Jovelin, and A. Dey, 2013 Molecular hyperdiversity and evolution in very large populations. Mol. Ecol. 22: 2074–2095. https://doi.org/10.1111/mec.12281

Cutter, A. D., and B. A. Payseur, 2013 Genomic signatures of selection at linked sites: unifying the disparity among species. Nat. Rev. Genet. 14: 262–274. https://doi.org/10.1038/nrg3425

Cutter, A. D., L. T. Morran, and P. C. Phillips, 2019 Males, Outcrossing, and Sexual Selection in Caenorhabditis Nematodes. Genetics 213: 27–57.

Danecek, P., A. Auton, G. Abecasis, C. A. Albers, E. Banks, et al., 2011 The variant call format and VCFtools. Bioinformatics 27: 2156–2158. https://doi.org/10.1093/bioinformatics/btr330

Dey, A., C. K. W. Chan, C. G. Thomas, and A. D. Cutter, 2013 Molecular hyperdiversity defines populations of the nematode Caenorhabditis brenneri. Proc. Natl. Acad. Sci. U. S. A. 110: 11056–11060. https://doi.org/10.1073/pnas.1303057110

Eyre-Walker A., 2006 The genomic rate of adaptive evolution. Trends Ecol. Evol. 21: 569–575. https://doi.org/10.1016/j.tree.2006.06.015

Eyre-Walker A., and P. D. Keightley, 2007 The distribution of fitness effects of new mutations. Nat. Rev. Genet. 8: 610–618. https://doi.org/10.1038/nrg2146

Eyre-Walker A., and P. D. Keightley, 2009 Estimating the rate of adaptive molecular evolution in the presence of slightly deleterious mutations and population size change. Molecular Biology and Evolution 26: 2097–2108.

Eyre-Walker A., 2010 Evolution in health and medicine Sackler colloquium: Genetic architecture of a complex trait and its implications for fitness and genome-wide association studies. Proc. Natl. Acad. Sci. U. S. A. 107 Suppl 1: 1752–1756. https://doi.org/10.1073/pnas.0906182107

Haller, B. C., and P. W. Messer, 2019 SLiM 3: Forward Genetic Simulations Beyond the Wright–Fisher Model. Molecular Biology and Evolution 36: 632–637.

Hill, W. G., and A. Robertson, 1966 The effect of linkage on limits to artificial selection. Genet. Res. 8: 269–294.

Huber, C. D., A. Durvasula, A. M. Hancock, and K. E. Lohmueller, 2018 Gene expression drives the evolution of dominance. Nat. Commun. 9: 2750. https://doi.org/10.1038/s41467-018-05281-7

Keightley, P. D., and A. Eyre-Walker, 2007 Joint inference of the distribution of fitness effects of deleterious mutations and population demography based on nucleotide polymorphism frequencies. Genetics 177: 2251–2261. https://doi.org/10.1534/genetics.107.080663

Kim, B. Y., C. D. Huber, and K. E. Lohmueller, 2017 Inference of the Distribution of Selection Coefficients for New Nonsynonymous Mutations Using Large Samples. Genetics 206: 345–361. https://doi.org/10.1534/genetics.116.197145

Kondrashov, A. S., 1995 Modifiers of mutation-selection balance: general approach and the evolution of mutation rates. Genetical Research 66: 53–69.

Kousathanas, A., and P. D. Keightley, 2013 A comparison of models to infer the distribution of fitness effects of new mutations. Genetics 193: 1197–1208. https://doi.org/10.1534/genetics.112.148023

Lee, R. Y. N., K. L. Howe, T. W. Harris, V. Arnaboldi, S. Cain, et al., 2018 WormBase 2017: molting into a new stage. Nucleic Acids Res. 46: D869–D874. https://doi.org/10.1093/nar/gkx998

Lee, D., S. Zdraljevic, L. Stevens, Y. Wang, R. E. Tanny, et al., 2020 Balancing selection maintains ancient genetic diversity in C. elegans. bioRxiv 2020.07.23.218420.

Li, H., 2011 A statistical framework for SNP calling, mutation discovery, association mapping and population genetical parameter estimation from sequencing data. Bioinformatics 27: 2987–2993. https://doi.org/10.1093/bioinformatics/btr509

Loewe, L., B. Charlesworth, C. Bartolomé, and V. Nöel, 2006 Estimating selection on nonsynonymous mutations. Genetics 172: 1079–1092. https://doi.org/10.1534/genetics.105.047217

Loewe, L., 2006 Quantifying the genomic decay paradox due to Muller’s ratchet in human mitochondrial DNA. Genet. Res. 87: 133–159. https://doi.org/10.1017/S0016672306008123

Lynch, M., 2008 The cellular, developmental and population-genetic determinants of mutation-rate evolution. Genetics 180: 933–943. https://doi.org/10.1534/genetics.108.090456

Messer, P. W., and D. A. Petrov, 2013 Population genomics of rapid adaptation by soft selective sweeps. Trends Ecol. Evol. 28: 659–669. https://doi.org/10.1016/j.tree.2013.08.003

Nordborg, M., 1997 Structured coalescent processes on different time scales. Genetics 146: 1501–1514.

Ohta, T., 1992 The Nearly Neutral Theory of Molecular Evolution. Annual Review of Ecology and Systematics 23: 263–286.

Peck, J. R., G. Barreau, and S. C. Heath, 1997 Imperfect genes, Fisherian mutation and the evolution of sex. Genetics 145: 1171–1199.

Pedersen, B. S., R. M. Layer, and A. R. Quinlan, 2016 Vcfanno: fast, flexible annotation of genetic variants. Genome Biol. 17: 118. https://doi.org/10.1186/s13059-016-0973-5

Pollak, E., 1987 On the theory of partially inbreeding finite populations. I. Partial selfing. Genetics 117: 353–360.

Poplin, R., V. Ruano-Rubio, M. A. DePristo, T. J. Fennell, M. O. Carneiro, et al., 2018 Scaling accurate genetic variant discovery to tens of thousands of samples. bioRxiv 201178.

Quinlan, A. R., and I. M. Hall, 2010 BEDTools: a flexible suite of utilities for comparing genomic features. Bioinformatics 26: 841–842. https://doi.org/10.1093/bioinformatics/btq033

Rockman, M. V., and L. Kruglyak, 2009a Recombinational landscape and population genomics of *Caenorhabditis elegans*. PLoS Genet. 5: e1000419. https://doi.org/10.1371/journal.pgen.1000419

Rockman, M. V., and L. Kruglyak, 2009b Recombinational landscape and population genomics of Caenorhabditis elegans. PLoS Genet. 5: e1000419. https://doi.org/10.1371/journal.pgen.1000419

Saxena, A. S., M. P. Salomon, C. Matsuba, S.-D. Yeh, and C. F. Baer, 2019 Evolution of the Mutational Process under Relaxed Selection inCaenorhabditis elegans. Molecular Biology and Evolution 36: 239–251.

Schultz, S. T., and M. Lynch, 1997 MUTATION AND EXTINCTION: THE ROLE OF VARIABLE MUTATIONAL EFFECTS, SYNERGISTIC EPISTASIS, BENEFICIAL MUTATIONS, AND DEGREE OF OUTCROSSING. Evolution 51: 1363–1371. https://doi.org/10.1111/j.1558-5646.1997.tb01459.x

Tataru, P., M. Mollion, S. Glémin, and T. Bataillon, 2017 Inference of Distribution of Fitness Effects and Proportion of Adaptive Substitutions from Polymorphism Data. Genetics 207: 1103–1119. https://doi.org/10.1534/genetics.117.300323

Thomas, C. G., W. Wang, R. Jovelin, R. Ghosh, T. Lomasko, et al., 2015 Full-genome evolutionary histories of selfing, splitting, and selection in Caenorhabditis. Genome Res. 25: 667–678. https://doi.org/10.1101/gr.187237.114

